# The genetic origin and architecture of sex determination

**DOI:** 10.1101/2021.12.06.471508

**Authors:** Tisham De

## Abstract

Here, I demonstrate that sex determination and sexual dimorphism across tree of life are deeply related to polyamine biochemistry in cells, especially to the synteny of genes: [SAT1-NR0B1], [SAT2-SHBG] and DMRT1. This synteny was found to be most distinct in mammals. Further, the common protein domain of SAT1 and SAT2 - PF00583 was shown to be present in the genome of the last universal common ancestor (LUCA). Protein domain-domain interaction analysis of LUCA’s genes suggests the possibility that LUCA had developed an immune defence against viruses. This domain-domain interaction analysis is the first scientific evidence indicating that viruses existed at least 3.5 billions years ago and probably co-existed with LUCA on early Hadean Earth.

---

Weiss et al 2016^1^ used phylogenetic approaches to infer the physiology and habitat of the last universal common ancestor (LUCA) and reported 355 genes and its protein domains. One of LUCA’s domain PF00583 maps to two human genes SAT1 and SAT2. Here, I demonstrate that in humans and all mammals, 1) SAT2 always occurs on autosomes and lies in very close proximity to SHBG (sex hormone binding globulin), 2) SAT1 always occurs on X chromosome and lies near NR0B1 (DAX1 protein) and 3) DMRT1 is always on a different but autosomal chromosome (**Figure 1, Supplementary figure 1**) (**Supplementary table 1, Supplementary table 2)**. This synteny or genetic linkage suggests evolutionarily converged co-regulation of genes. Further, SAT1 and NROB1 tend to be located at a fixed distance of ∼6-7 mega base pairs (Mbps) in the genomes of all mammals (Humans=6.5 Mbps, Horse=5.6 Mbps, Blue Whale=6.9 Mbps), indicating that they have conserved long range interactions. SAT2 and SHBG on the other hand are usually located quite close to each other but with opposite orientation, suggesting transcriptional co-regulation. Some notable exceptions to these broad patterns were Petromyzon marinus (sea lamprey), Asterias rubens (common starfish) and some birds and animals endemic to the Australian continent.

**Figure 1.**
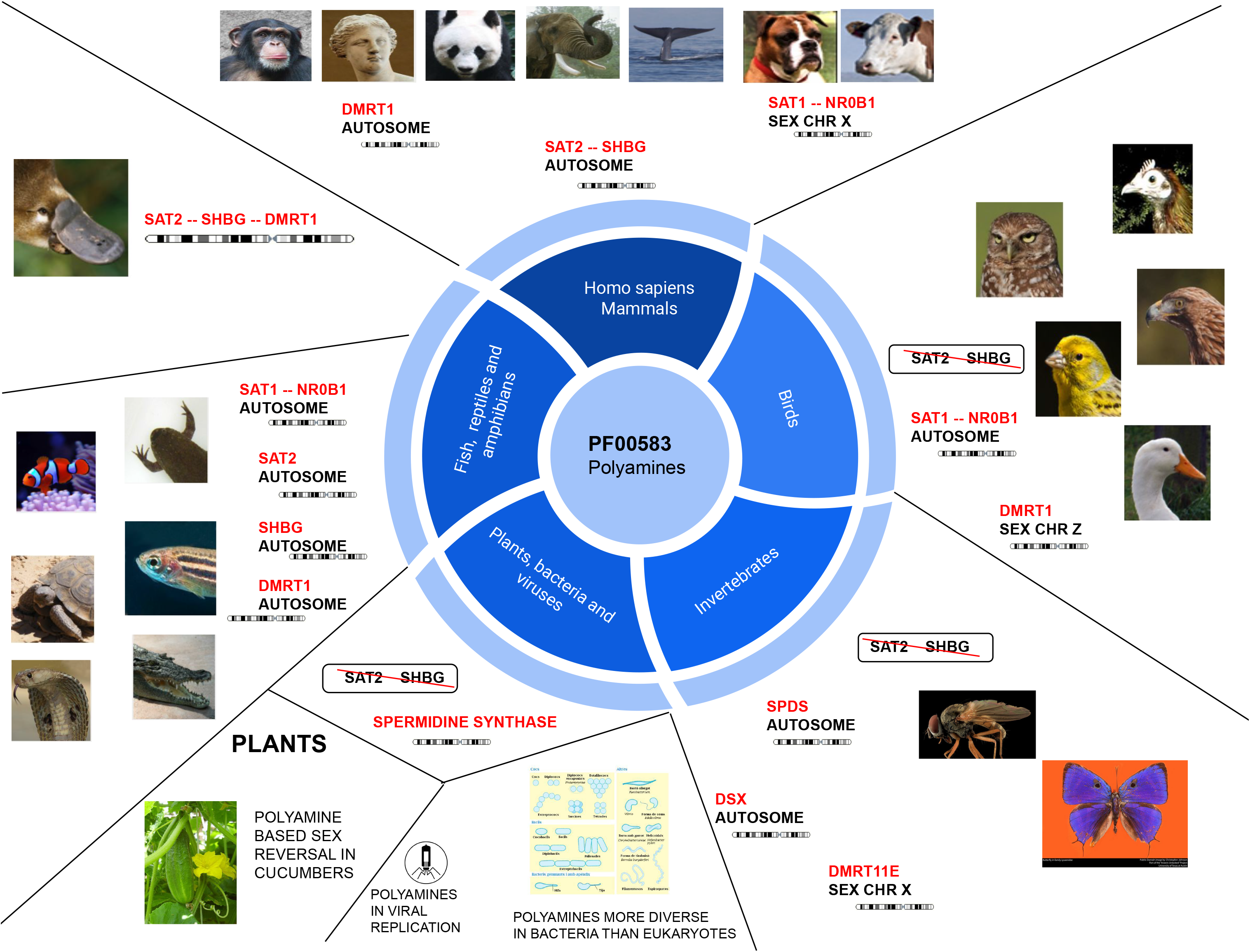
New taxonomy of tree of life based on polyamines.

Of note, in almost all human cancers the synteny of somatic structural variations of these genes is highly correlated and conserved (**Figure 2, Supplementary figure 2**), thus confirming and validating the common evolutionary trajectories observed in different species. The fact that this correlation in cancer genomes is mainly seen as structural variations (deletion and duplications) and not as somatic point mutations^2^ strongly indicates that the function of these genes and its effect on cellular phenotype (e.g. sex) is based on gene dosage and the concentrations of its protein products. These correlated somatic mutations perhaps define unique cellular states based on metabolism and homeostatis. Gene expression analysis from public databases ^3, 4^ further shows that SAT1 or spermidine biosynthesis related genes and pathways are universally expressed in almost all cell types (**Supplementary video 1**) across all species, where as SAT2, SHBG, NR0B1 and DMRT1 are very tissues specific, and likely to be influenced by other factors not known at the moment. Of note, chromosome Xp21.2-Xp22.1 duplications containing SAT1 and NR0B1 was earlier reported in sex reversed human individuals (male to female with normal 46, XY karyotype)^5–7^. Polyamines were also found to be involved in ethrel based sex reveral of cucumbers (Cucumis sativus L.)^8^. Based on current literature on polyamines (SAT1 and SAT2), NR0B1, SHBG and DMRT1, it appears that these genes are functionally connected through their common role in male testis formation by sertoli cells ^9, 10^. In birds, SAT1 and NR0B1 are always autosomal and co-located, SAT2 and SHBG are absent from the genome and DMRT1 always lie on the sex chromosome Z. In fish (which may not have distinct sex detemination system), amphibians and reptiles, most of these genes including SHBG are present (unlike birds), but with depleted events of co-location and usually do not occur on the same chromosome. Platypus (Ornithorhynchus anatinus) is the only species where SAT2, SHBG and DMRT1 were found to be co-located close to each other on the same chromosome. Further, taking into account suggestive evidence from invertebrates^11, 12^, plants^13–18^, bacteria, archaea^19–21^ and viruses^22, 23^, one may conclude that polyamines and its regulation are likely to be a universal and important component of the polygenic architecture of sex determination and sexual dimorphism across tree of life on Earth. An explanation for such a fundamental effect of polyamines like spermidine is perhaps because of its high positive charge (+3) which makes it a vital interaction partner for negatively charged nucleotides like DNA and RNA in the nucleus of the cell ^24^.

**Figure 2.**
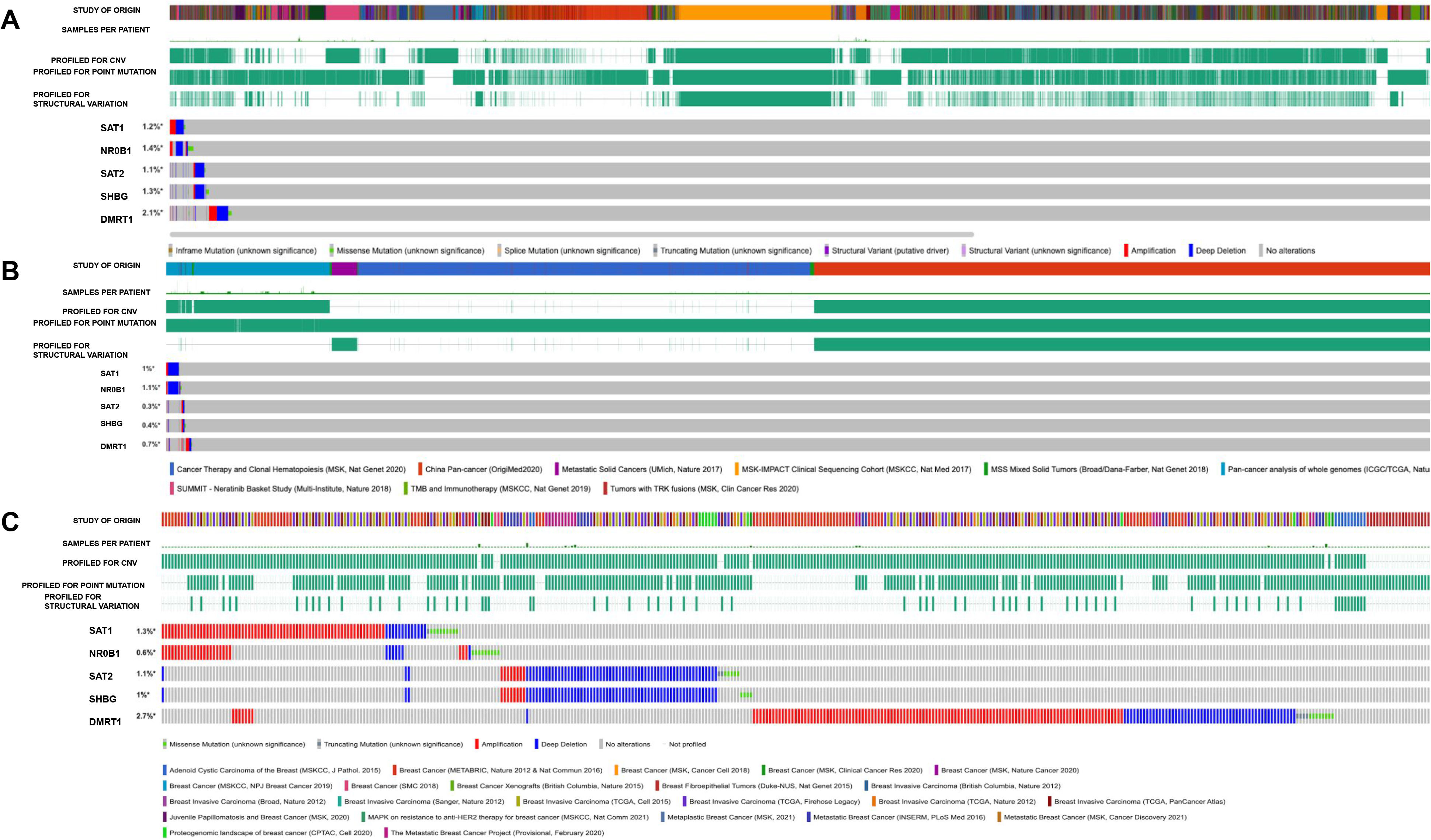
Synteny of [SAT1-NR0B1], [SAT2-SHBG] and DMRT1 in pan-cancer and breast cancer studies.

Protein domain-domain interaction analysis ^25, 26^ offers an alternative to protein-protein interaction analysis. Here, I have reported the protein domain-domain interaction analysis of the 355 domains reported by Weiss et al. 2016. Reactome^27^ pathway analysis (Homo sapiens) of the superset of protein domains obtained from this analysis shows strong enrichment for immune related genes and pathways. These observations indicate that LUCA’s protein domains and its interaction partners were perhaps part of larger biological pathways related to primitive immune defense against viruses (**Figure 3, Supplementary table 3**).

**Figure 3.**
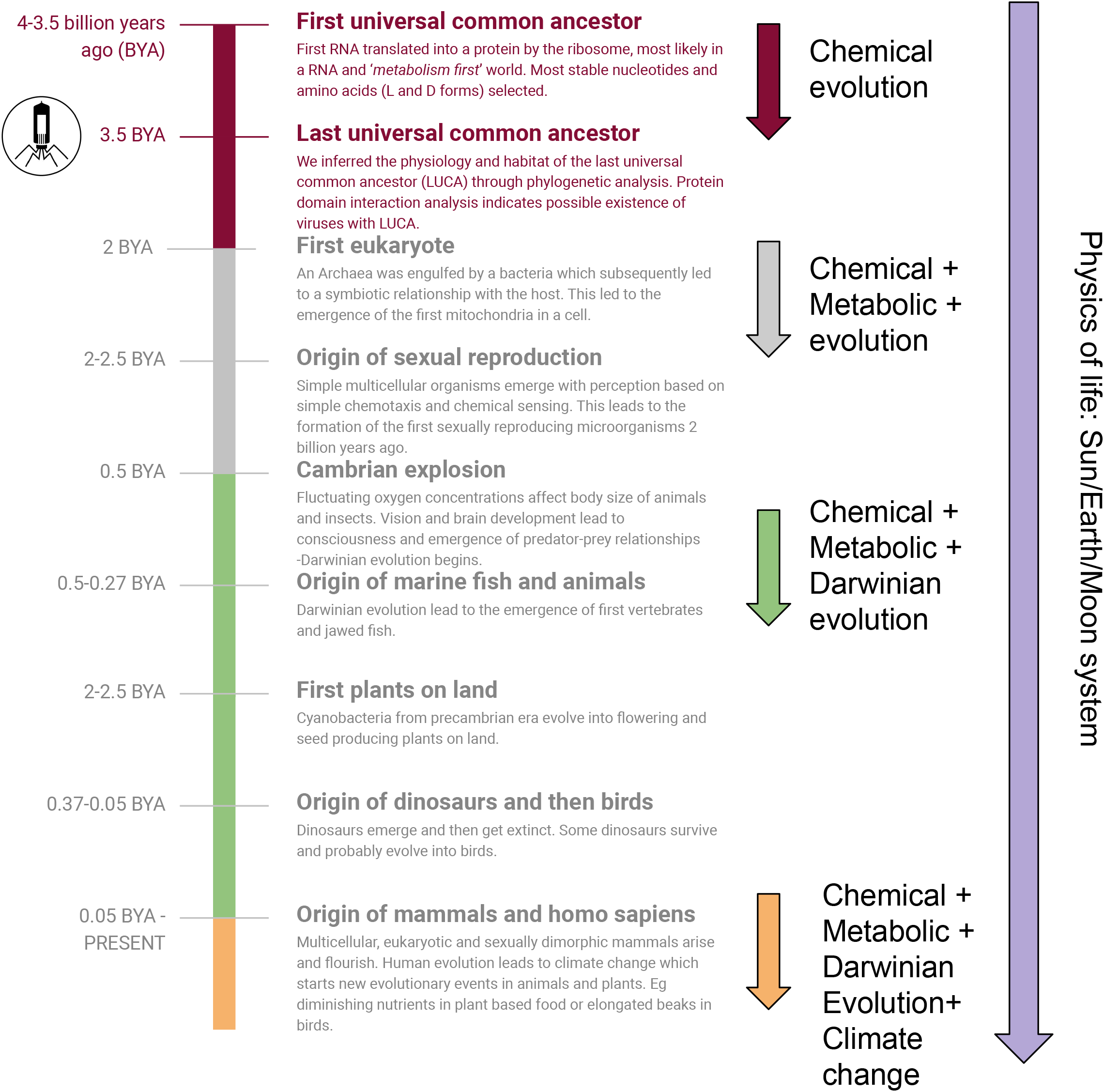
An overview of the evolution of life from LUCA to Homo Sapiens.

## AUTHOR CONTRIBUTIONS

**T.D** Involved in study design, performed analysis and wrote the manuscript.

## SUPPLEMENTAL TABLE TITLES

**Supplementary table 1.** Synteny results from UCSC genome browser

**Supplementary table 2.** Synteny results from the Ensembl genome browser.

**Supplementary table 3.** Reactome pathway results for LUCA’s domains and its interaction partners.

## SUPPLEMENTAL FIGURES

**Supplementary figure 1.** Gene expression results form GTEx

**Supplementary figure 2**. Synteny of [SAT1-NR0B1], [SAT2-SHBG] and DMRT1 in various cancer studies.

Supplementary video. Summary of single cell gene expression data (N=229, experiments) for SAT1 in various species. Url: https://www.ebi.ac.uk/gxa/sc/home

---

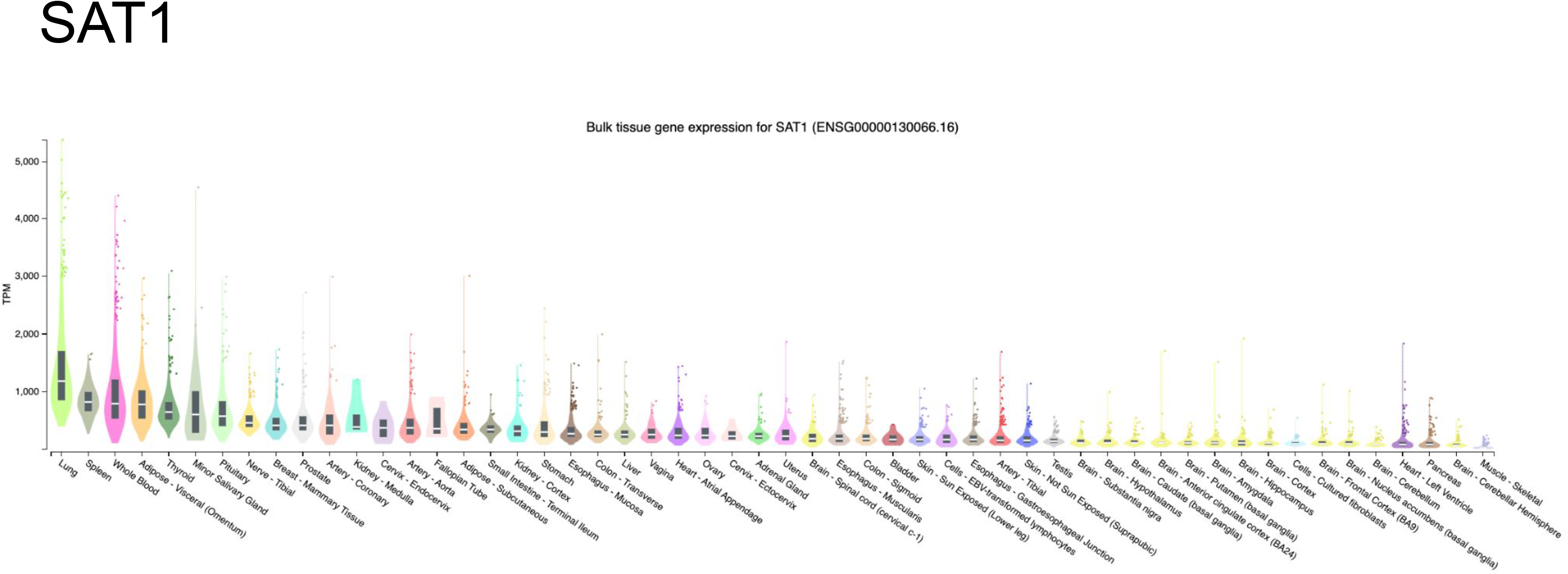
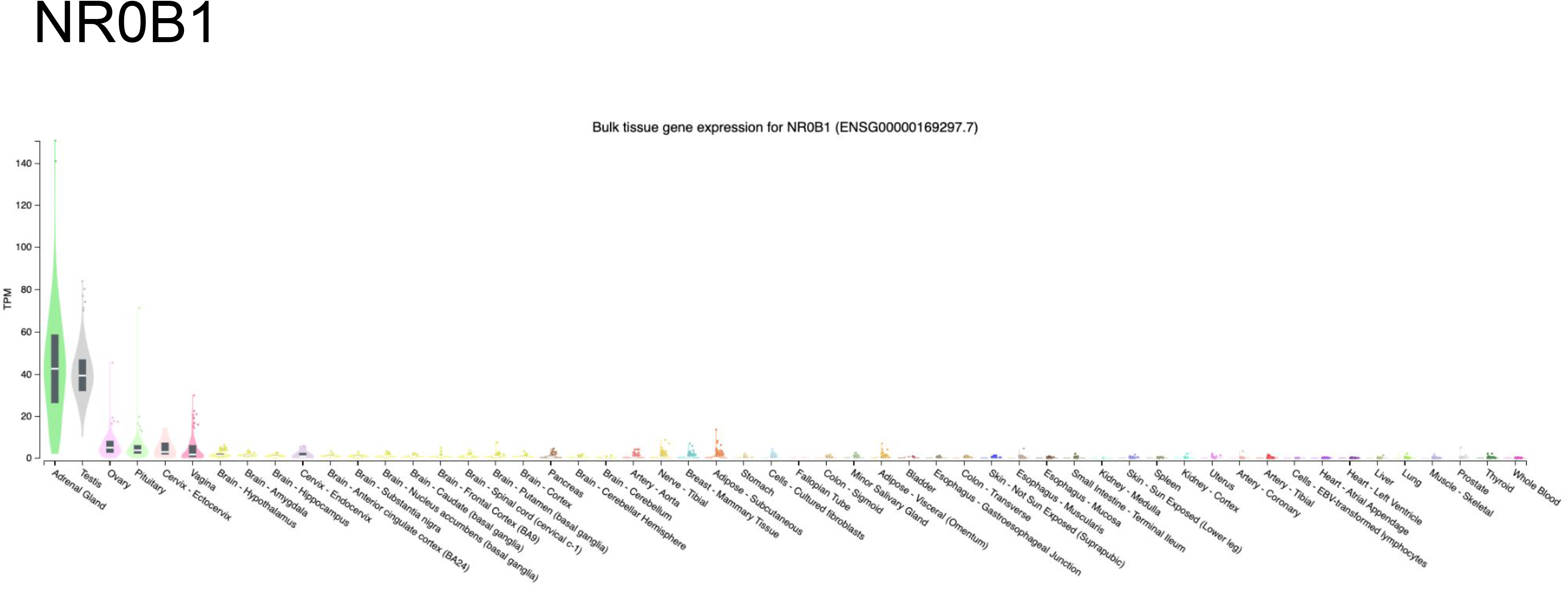
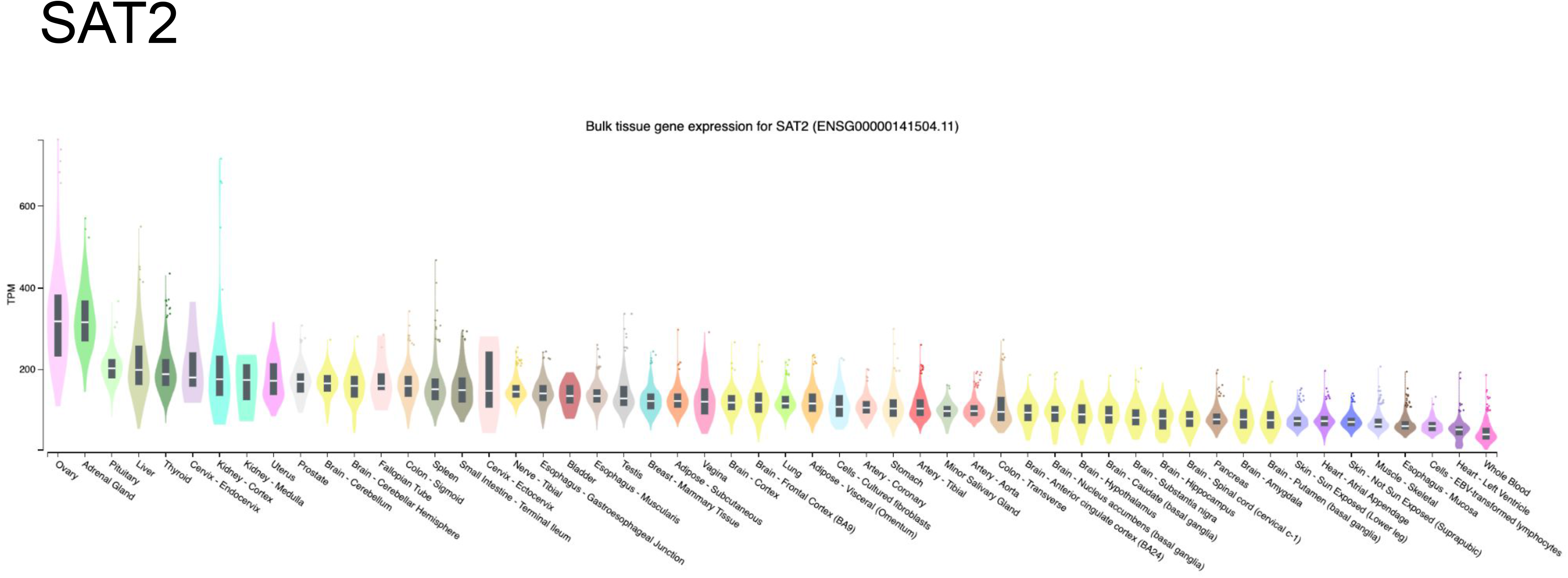
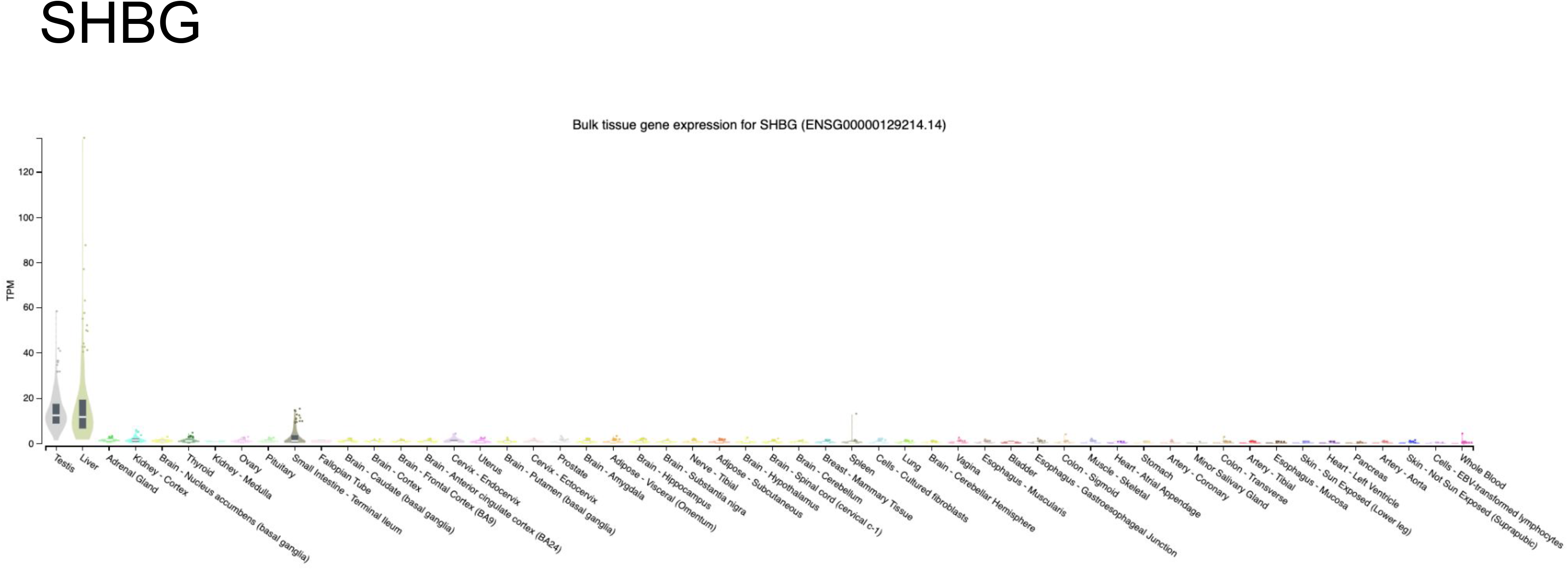
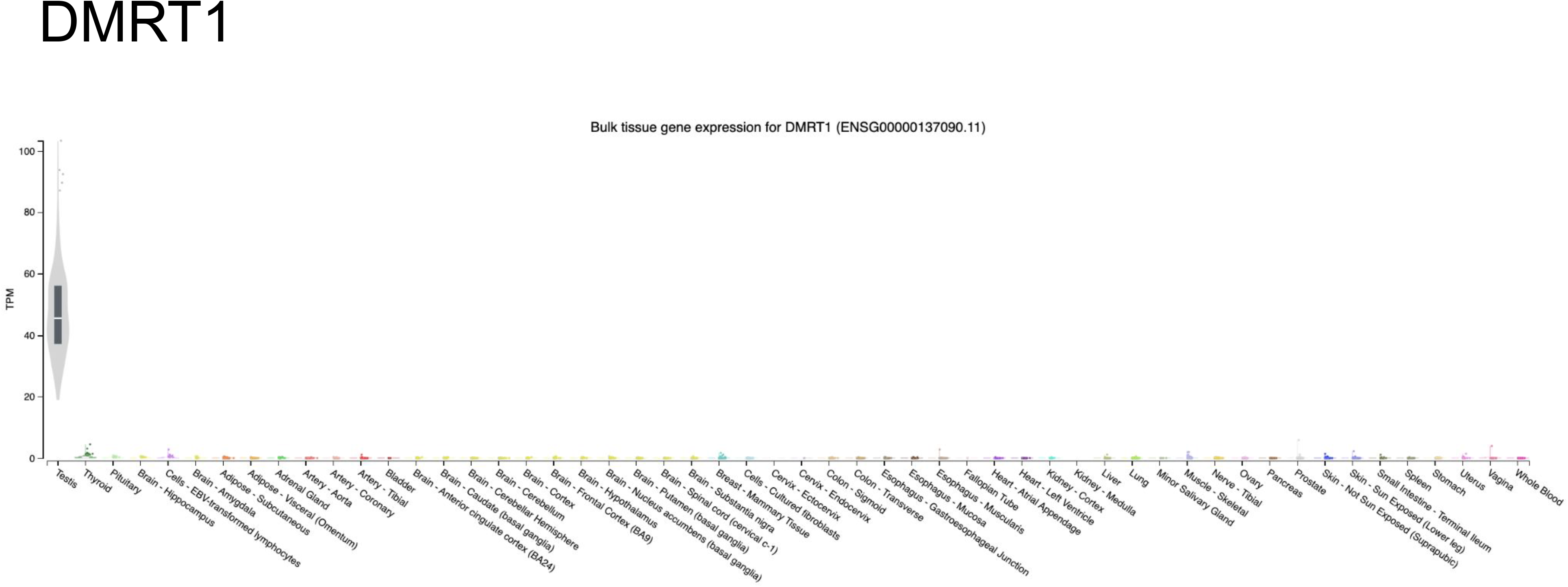
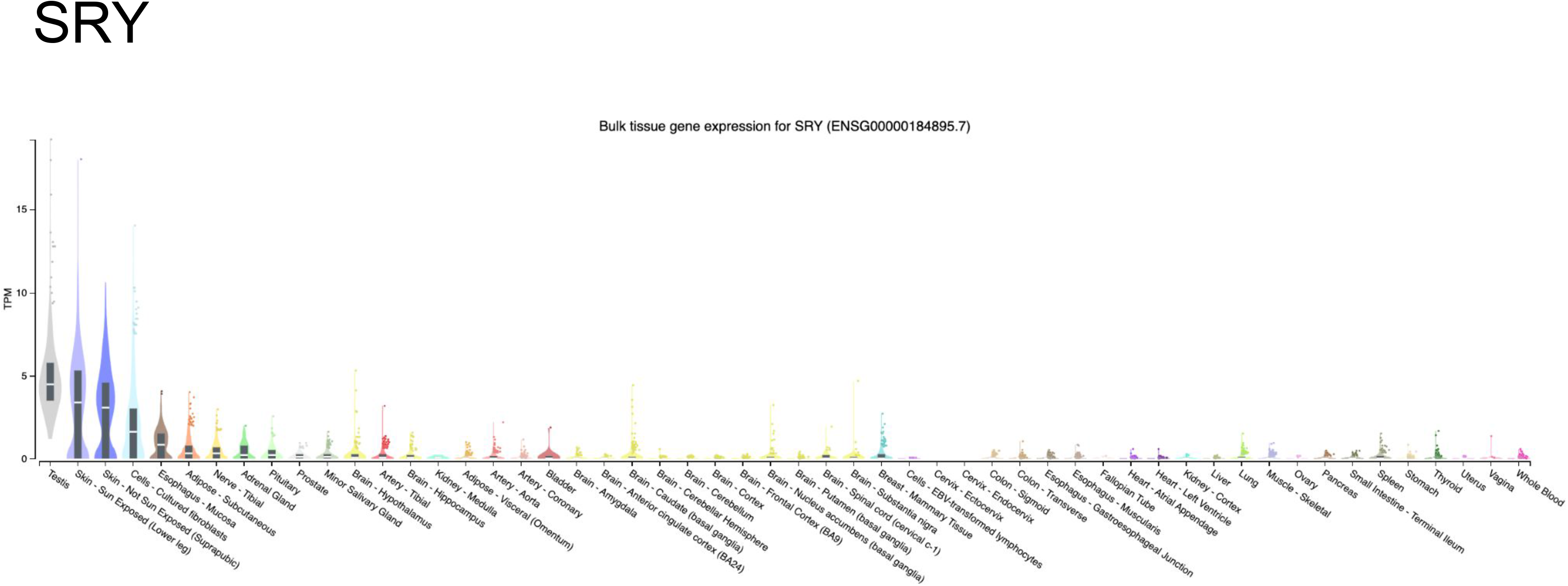
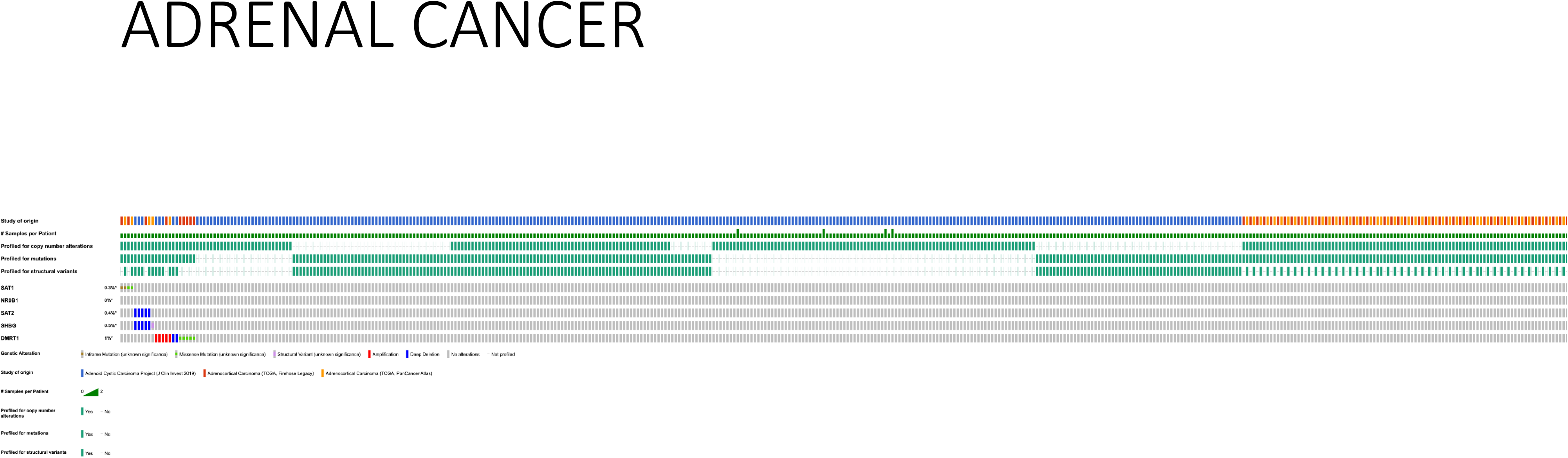
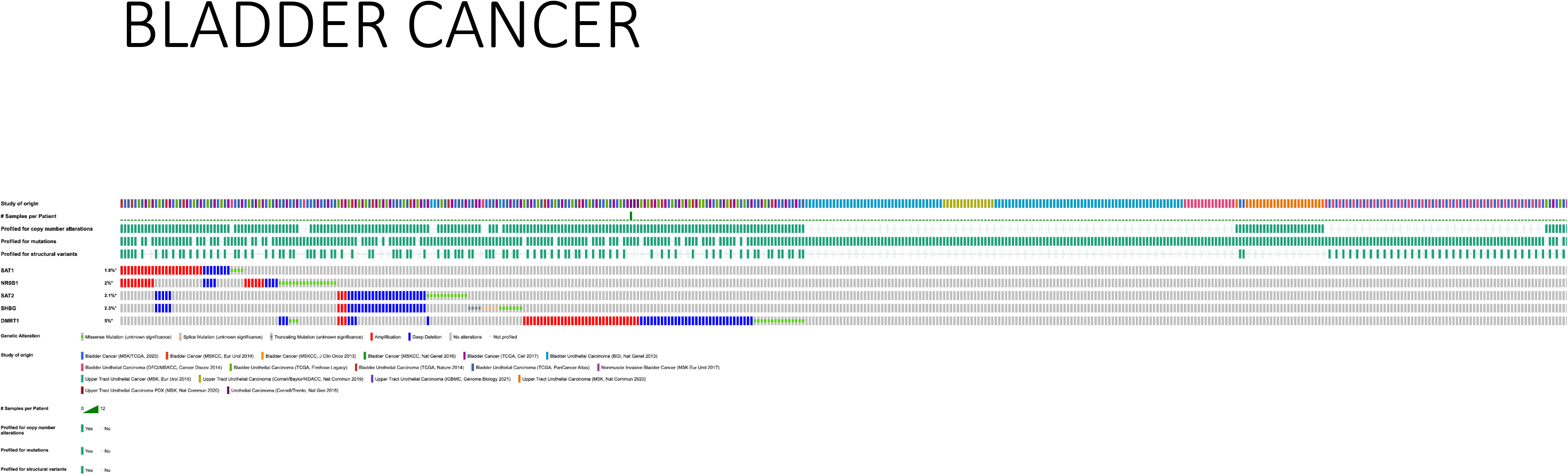
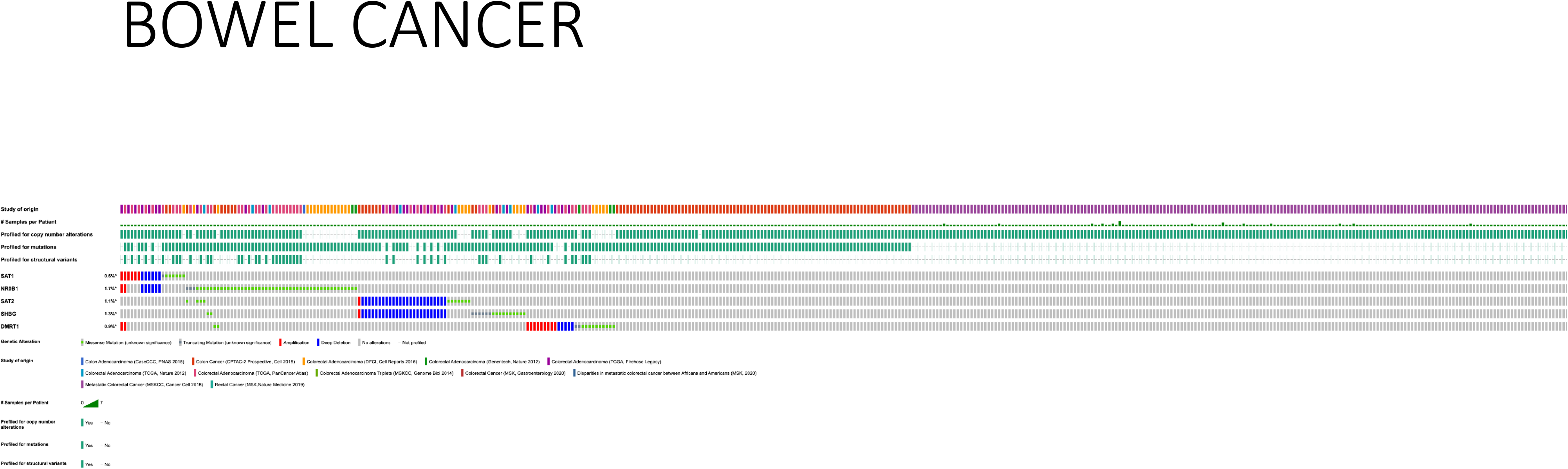
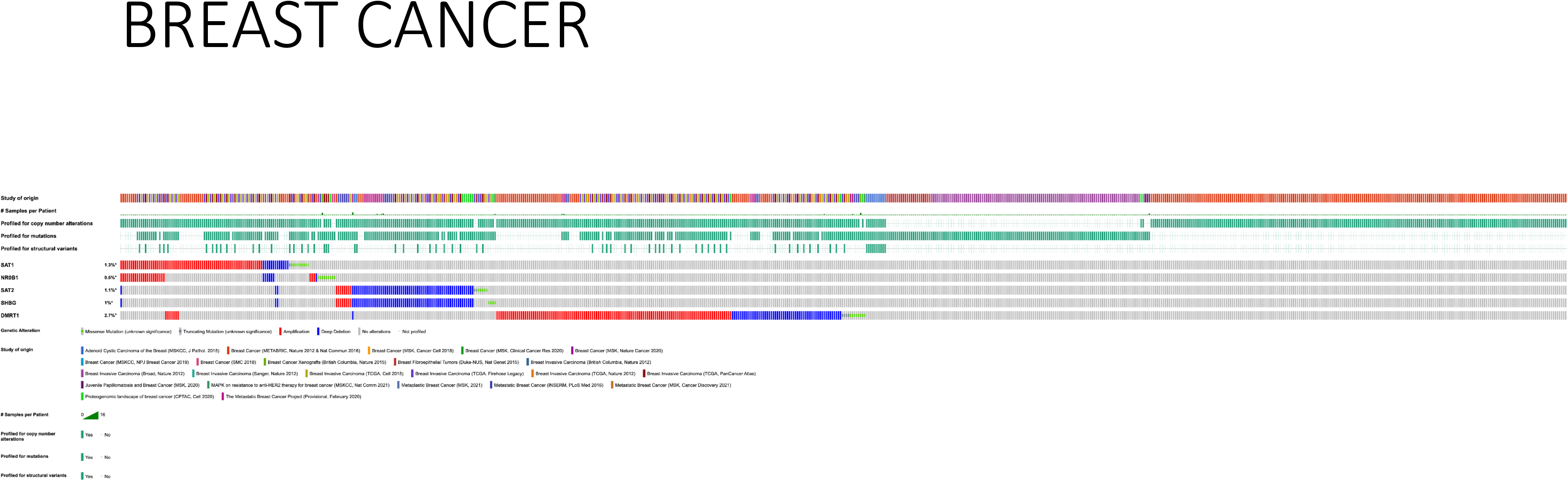
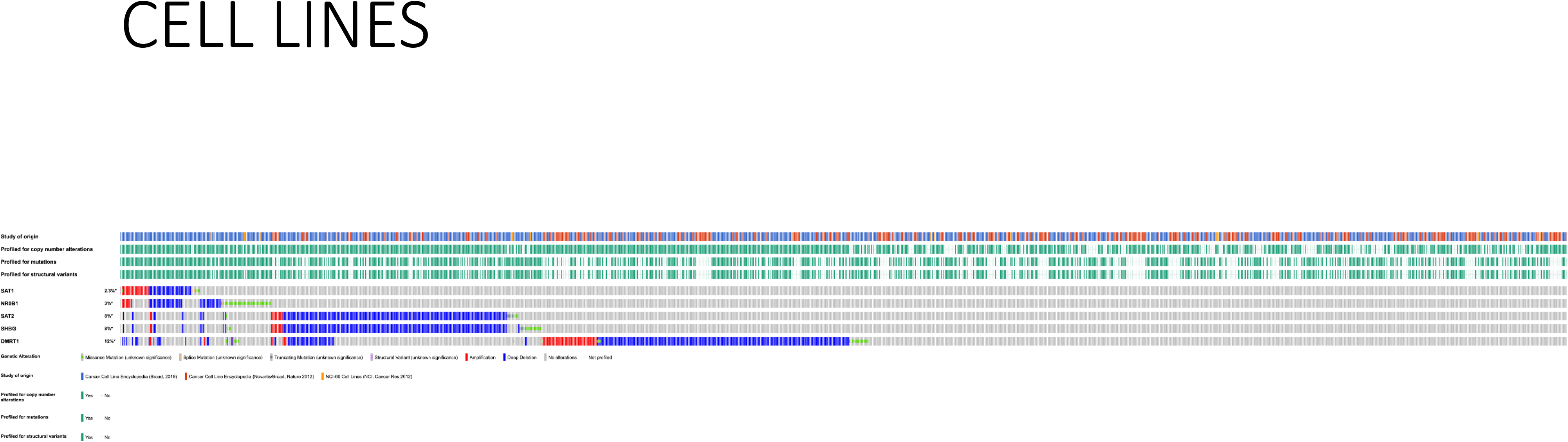
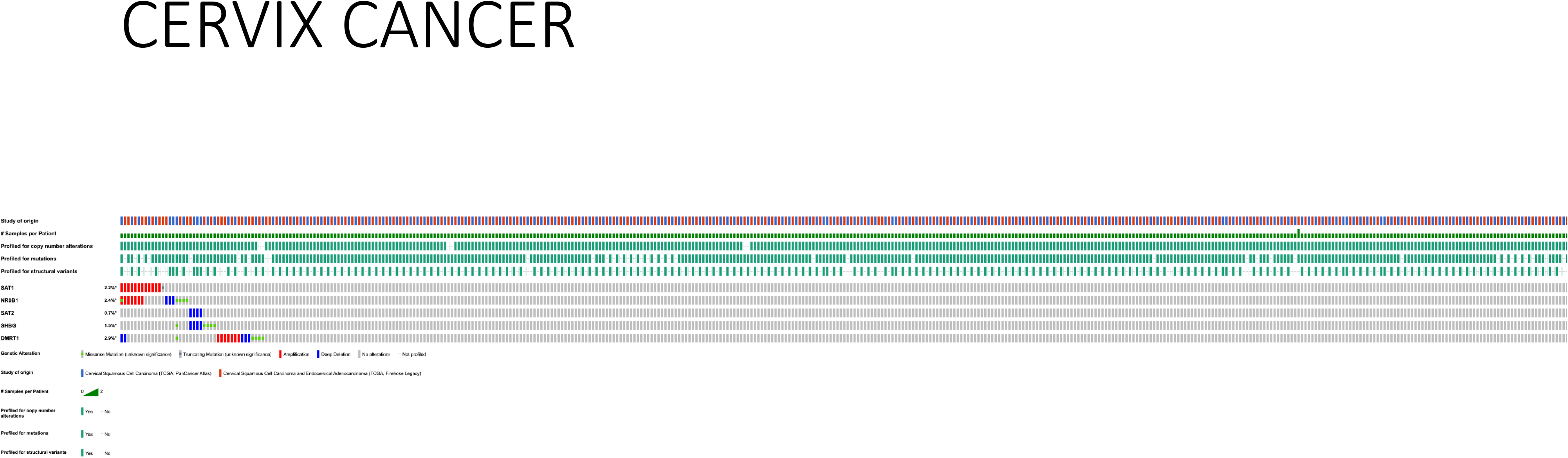
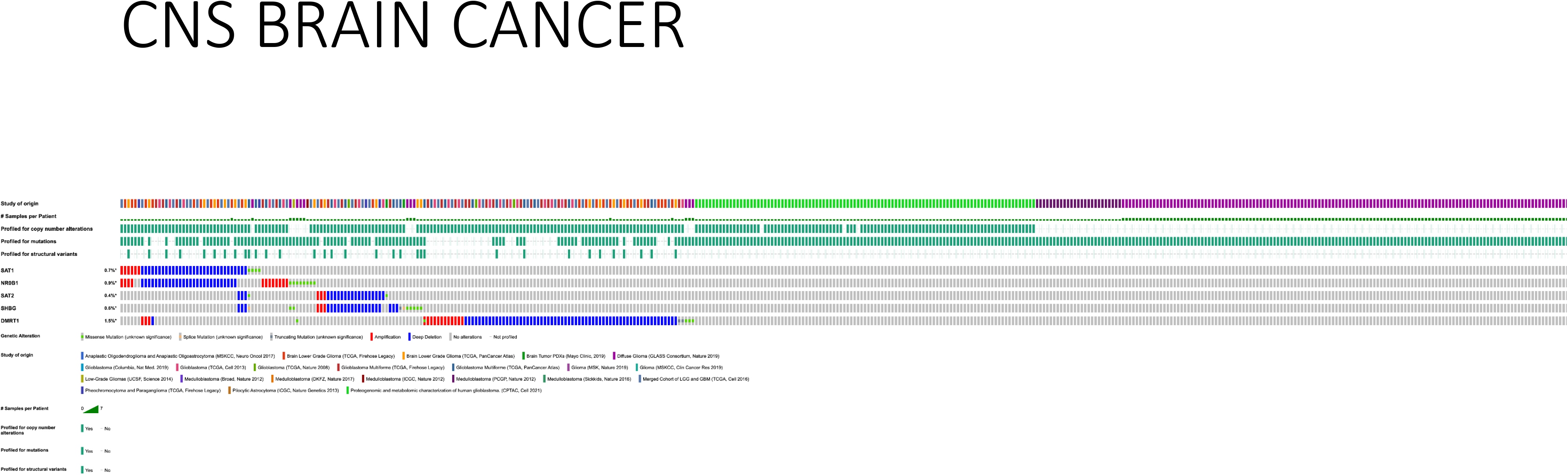
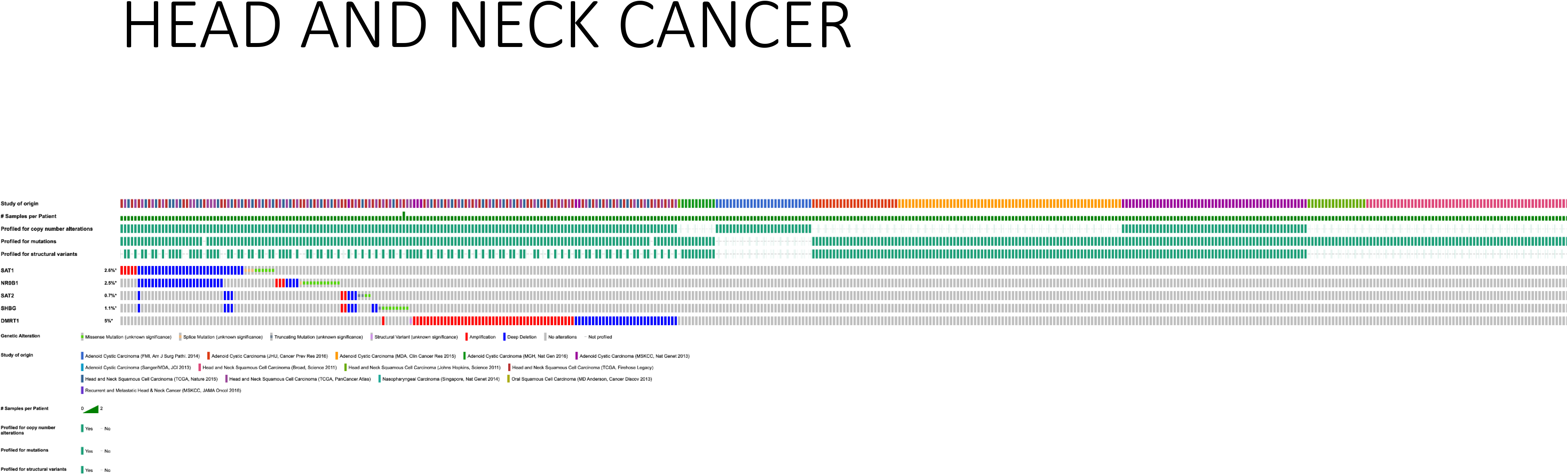
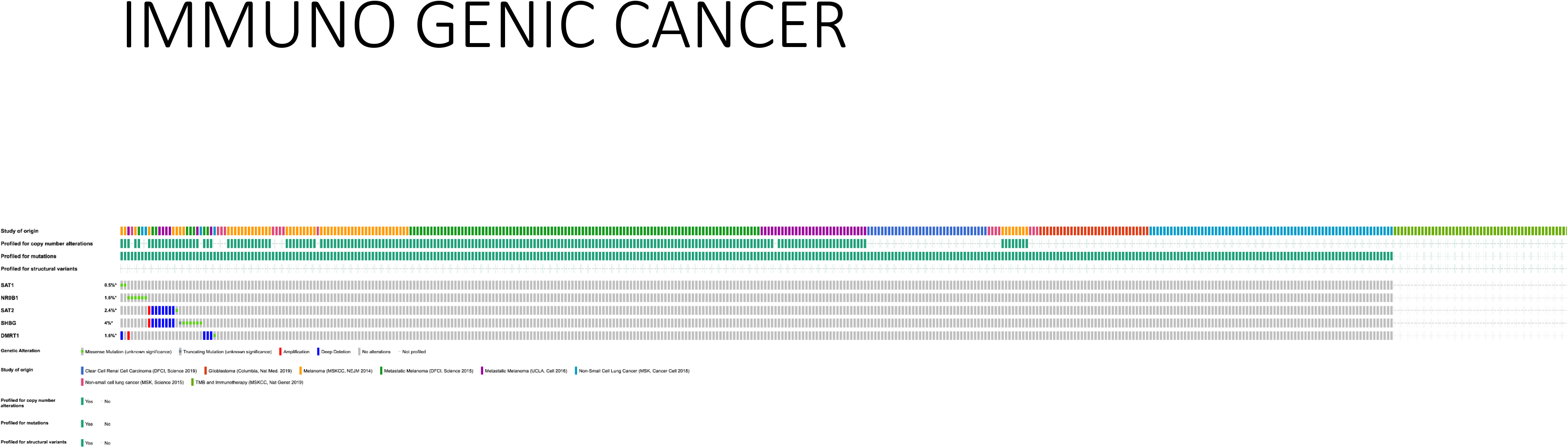
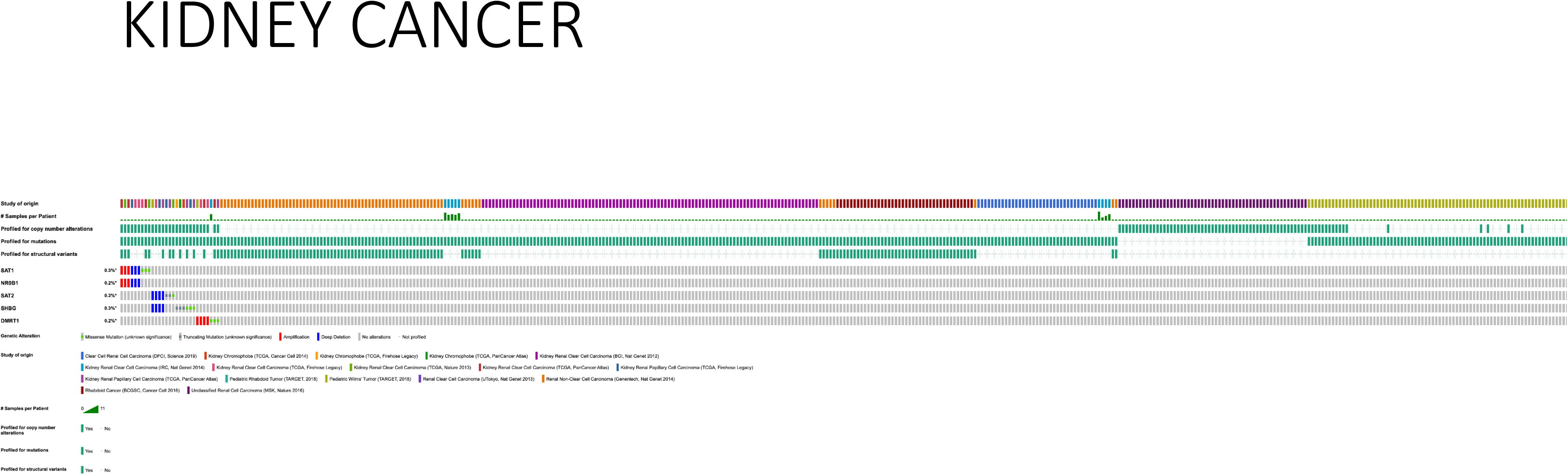
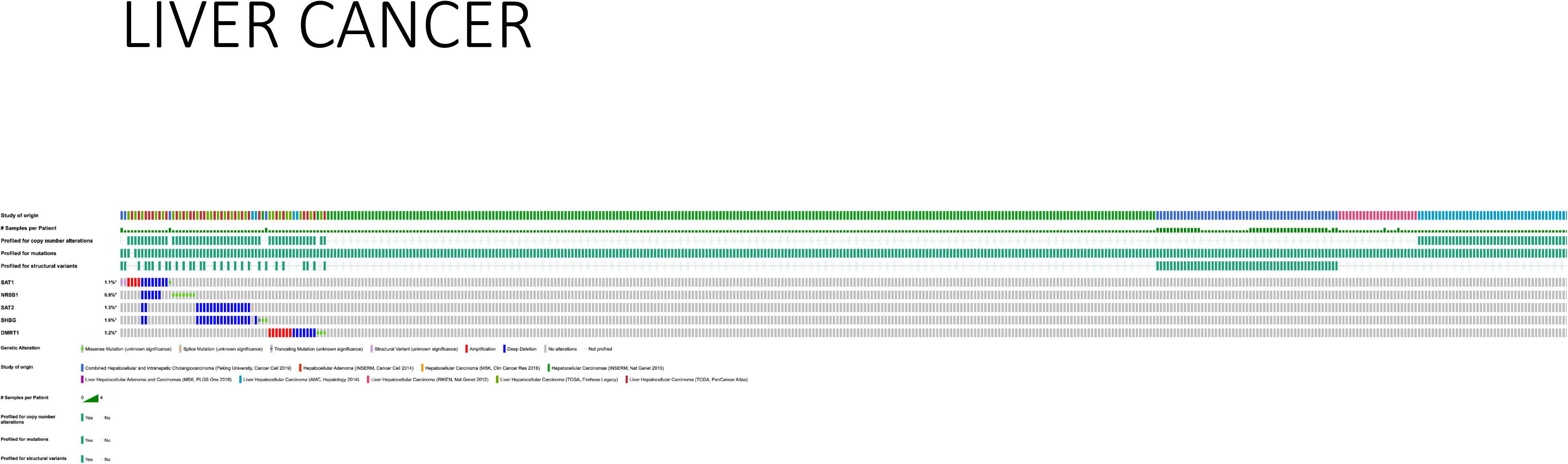
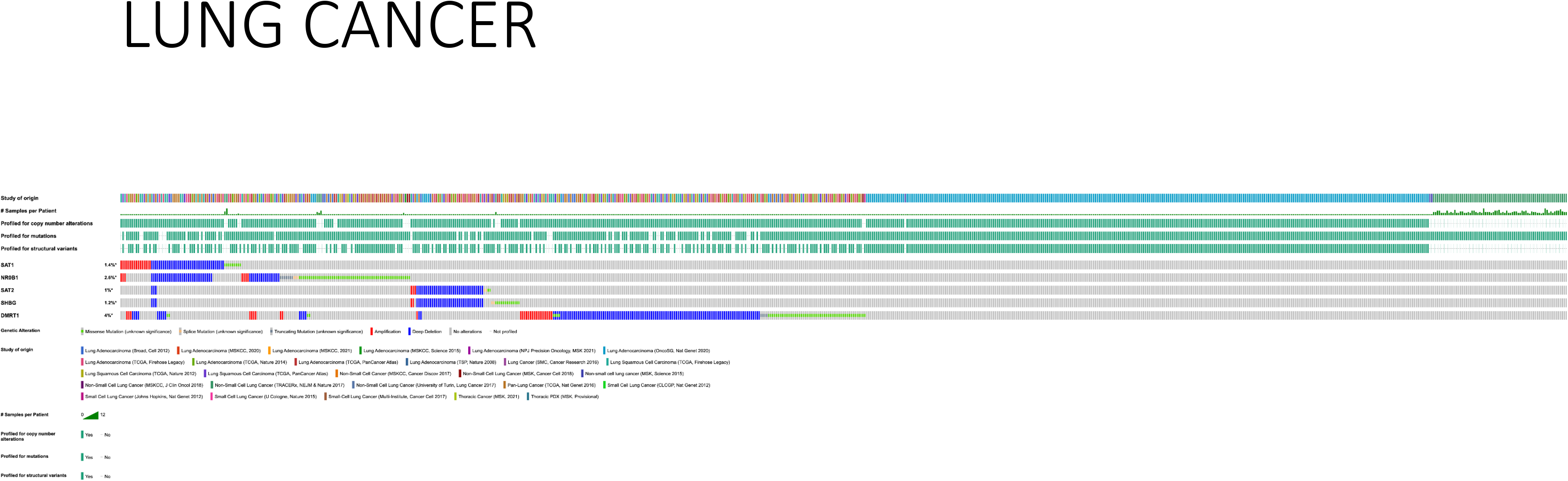
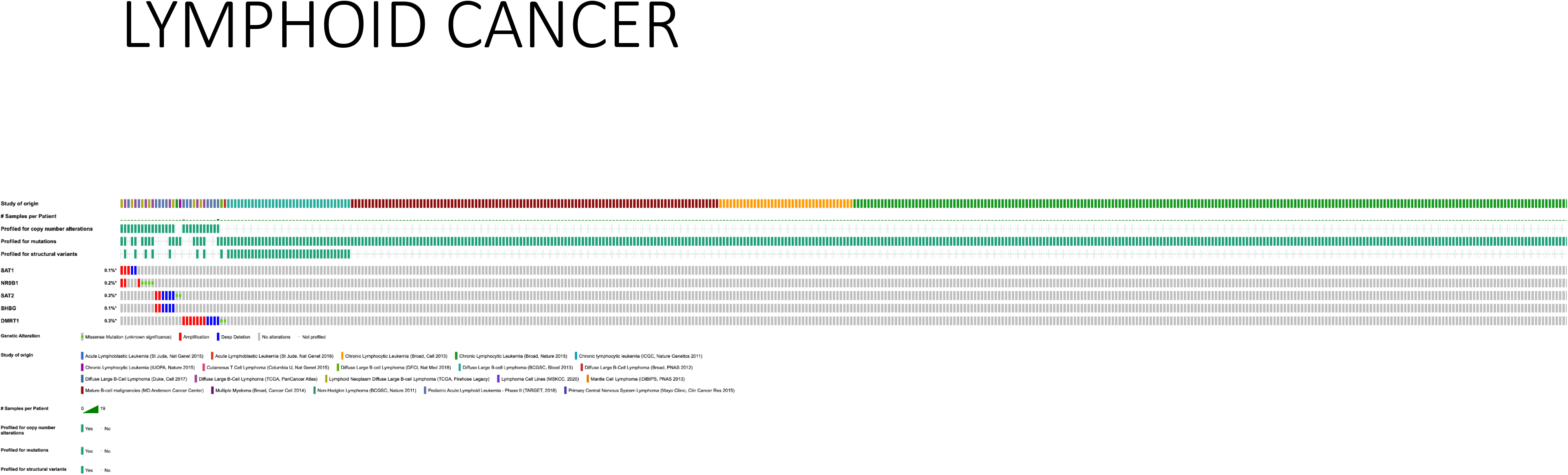
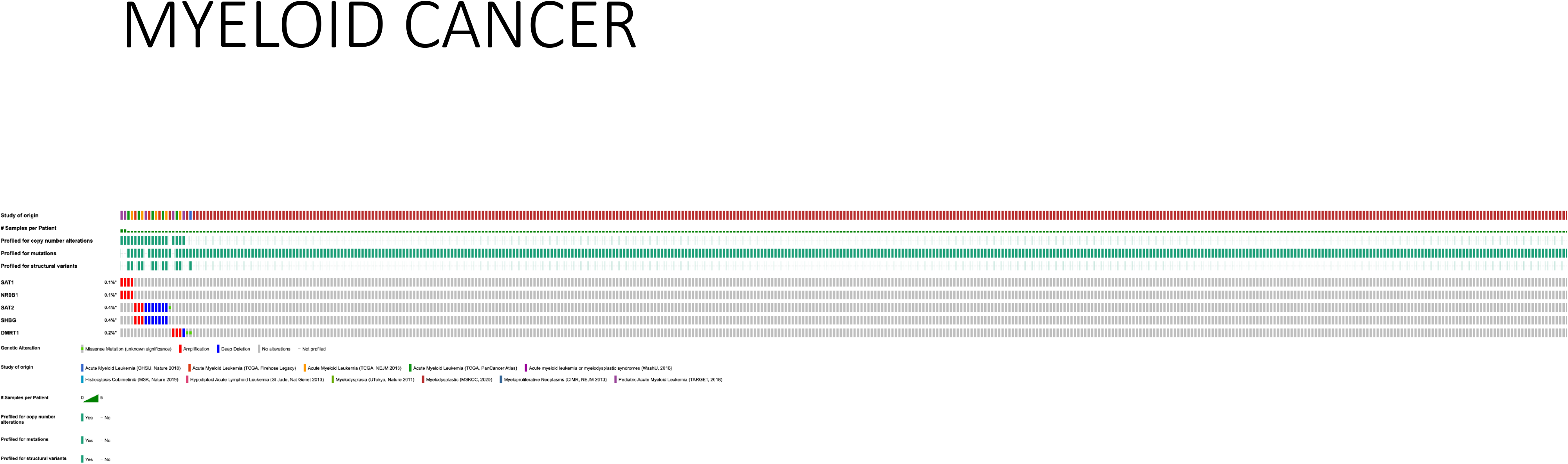
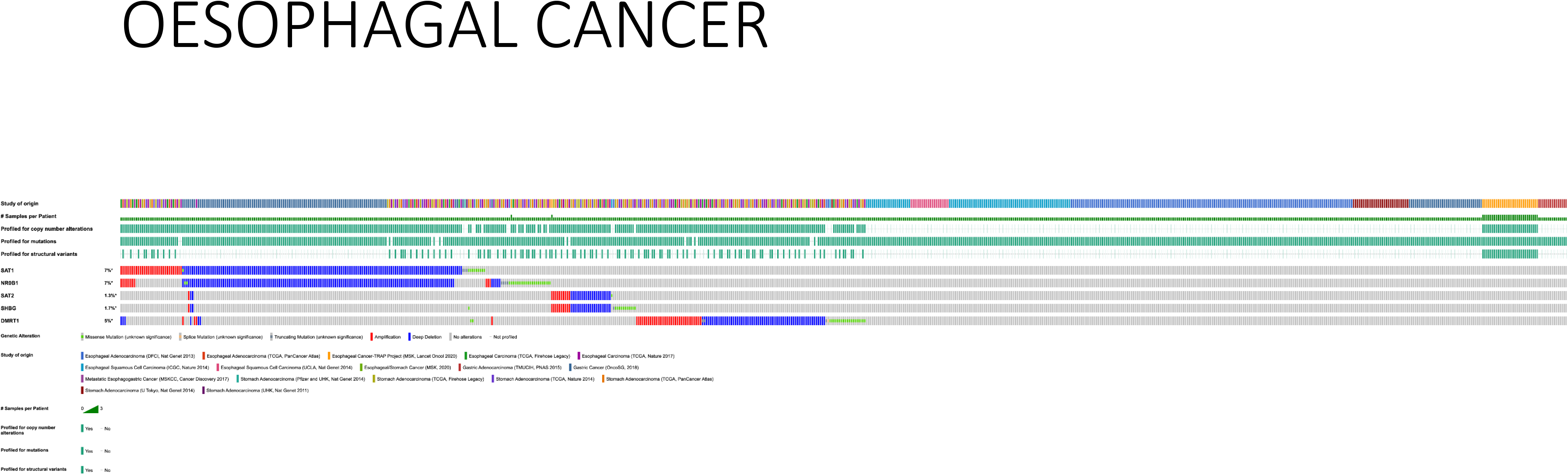
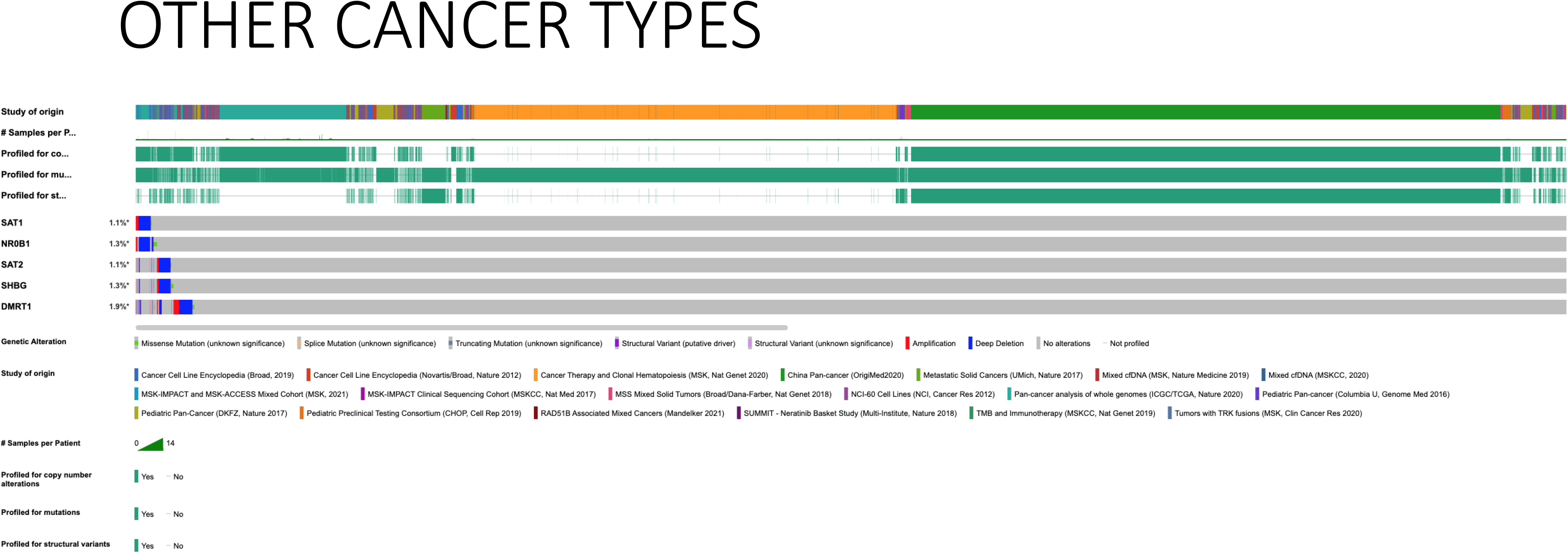
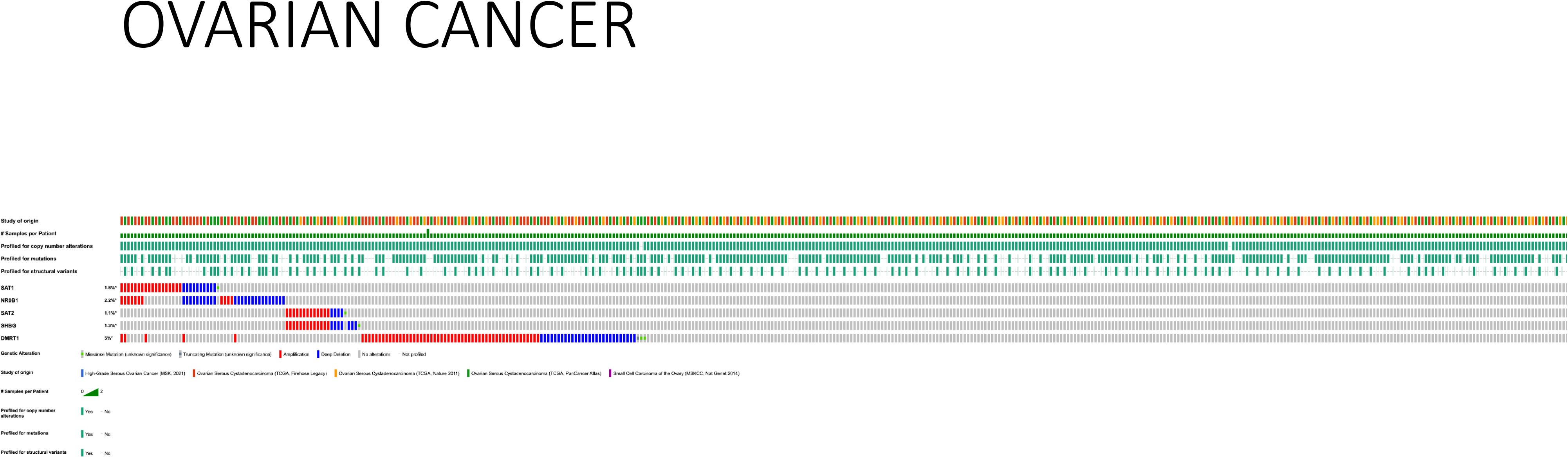
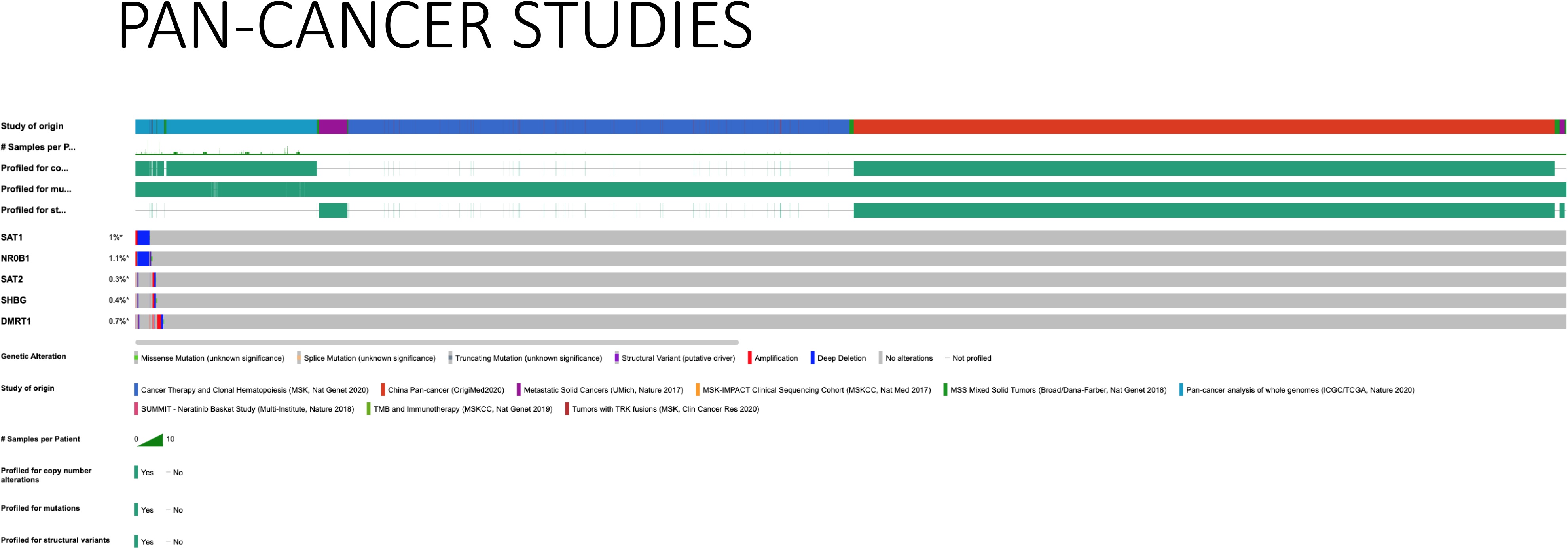
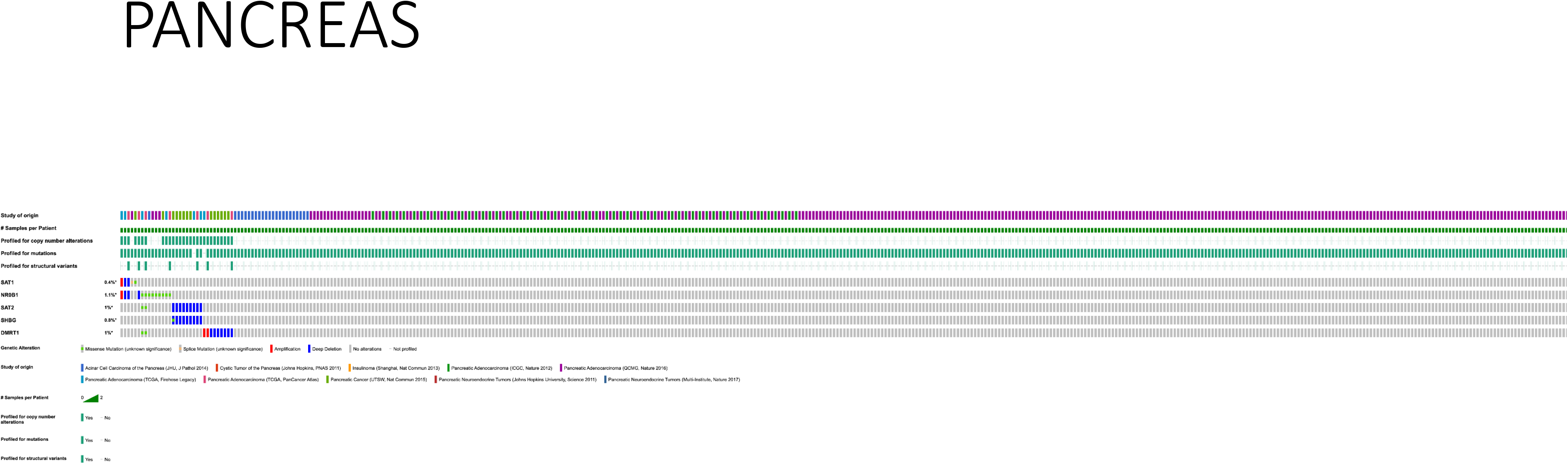
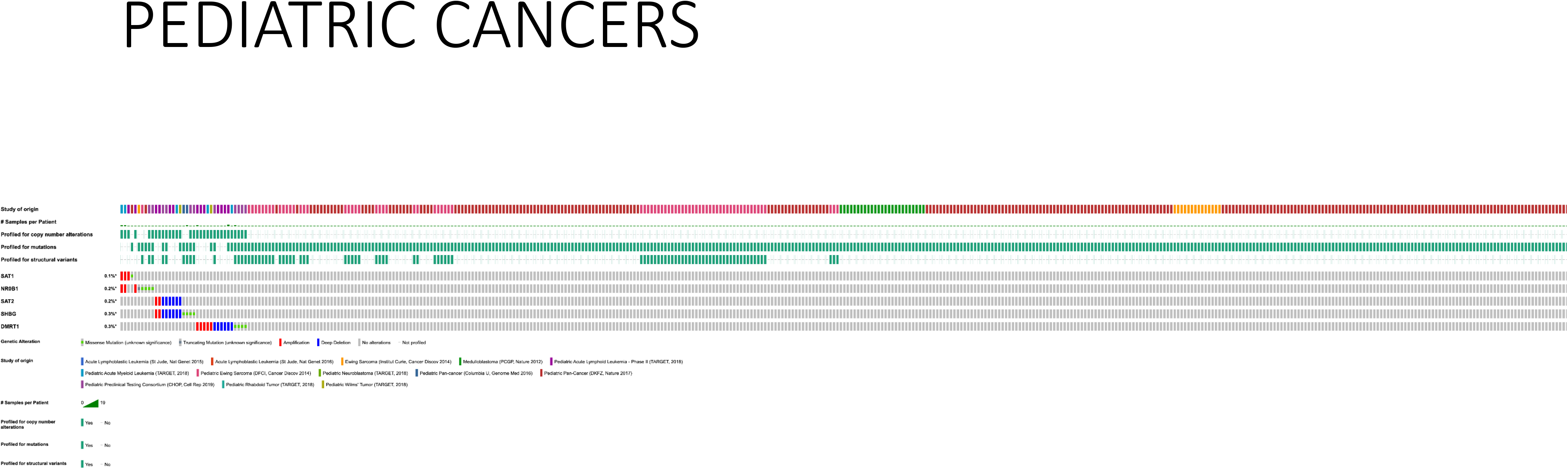
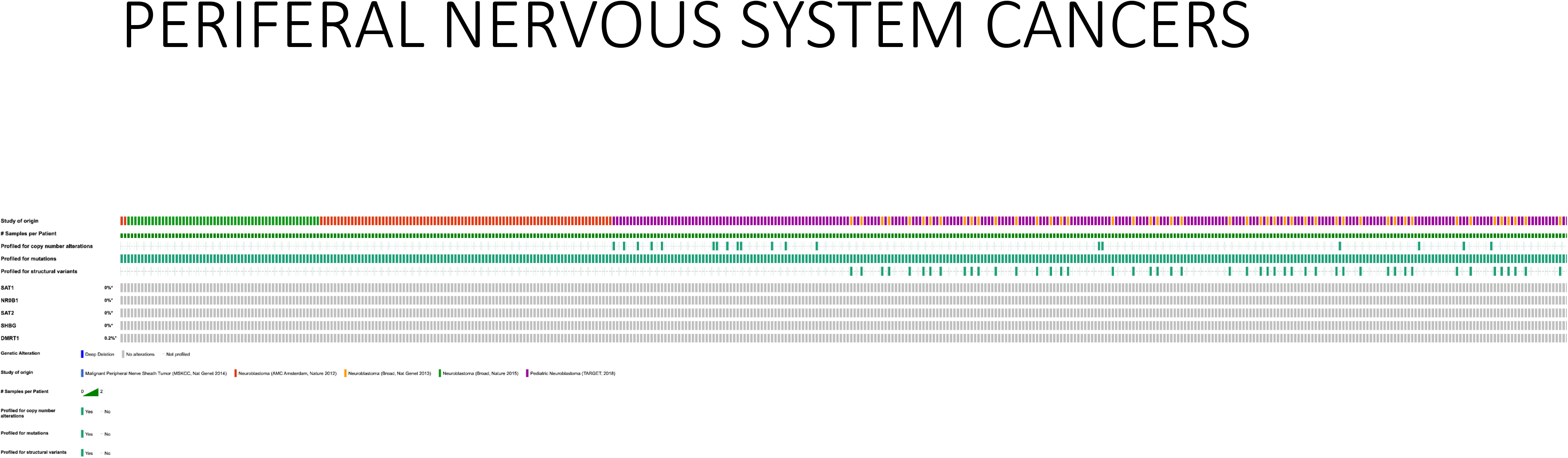
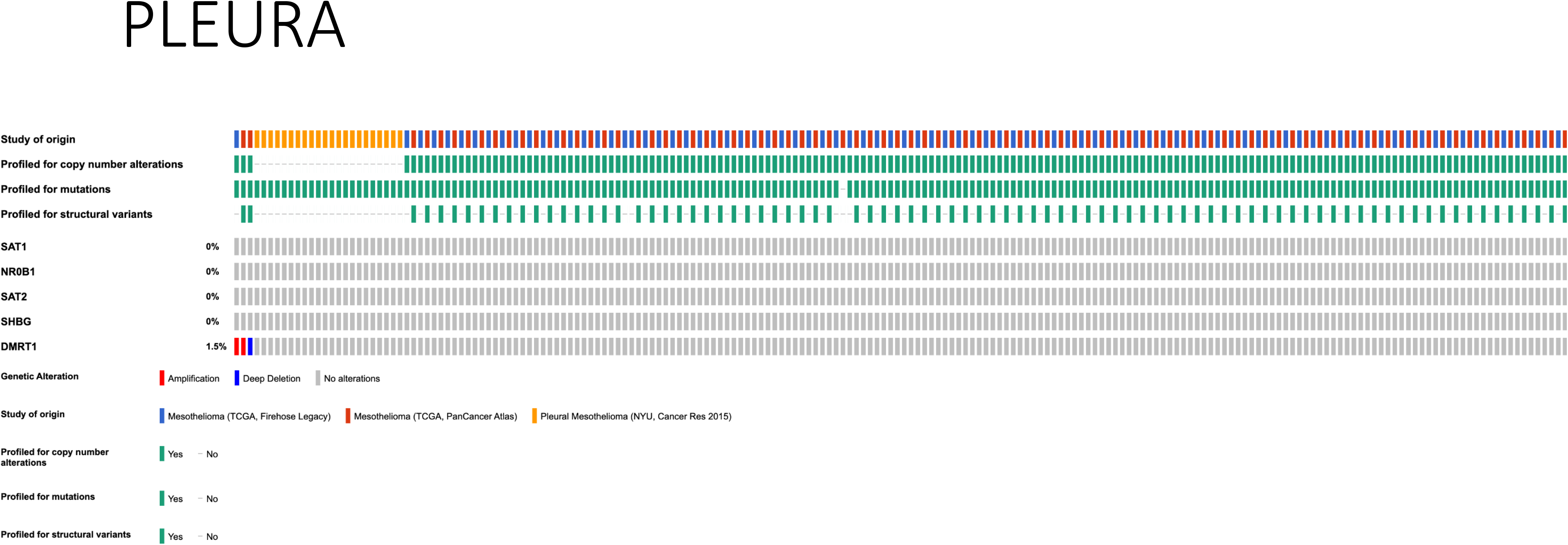
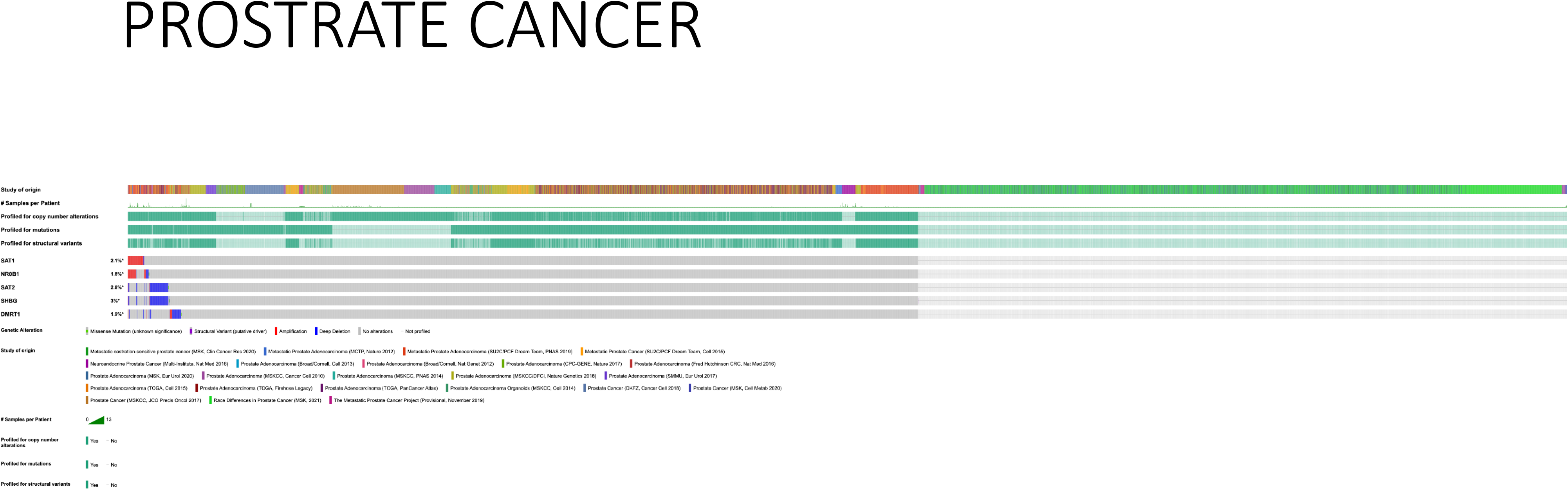
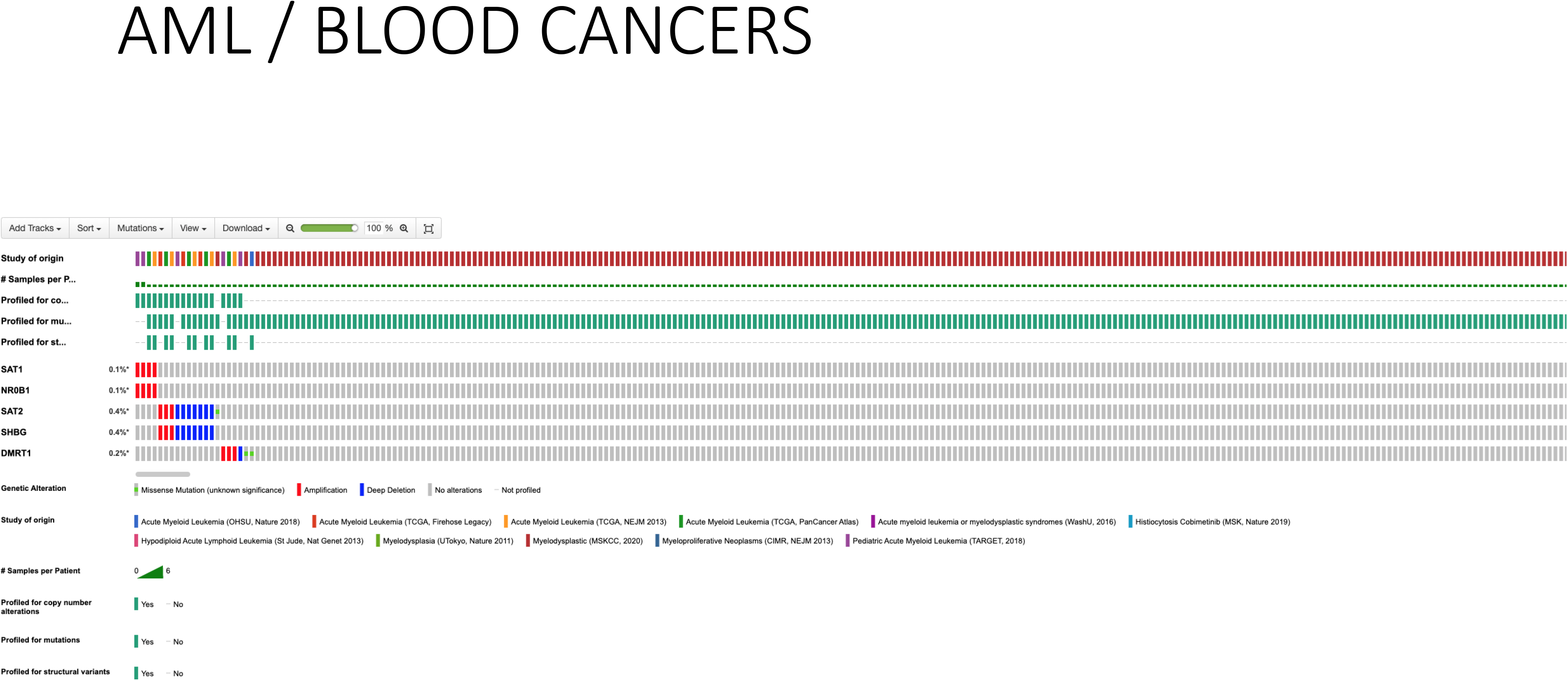
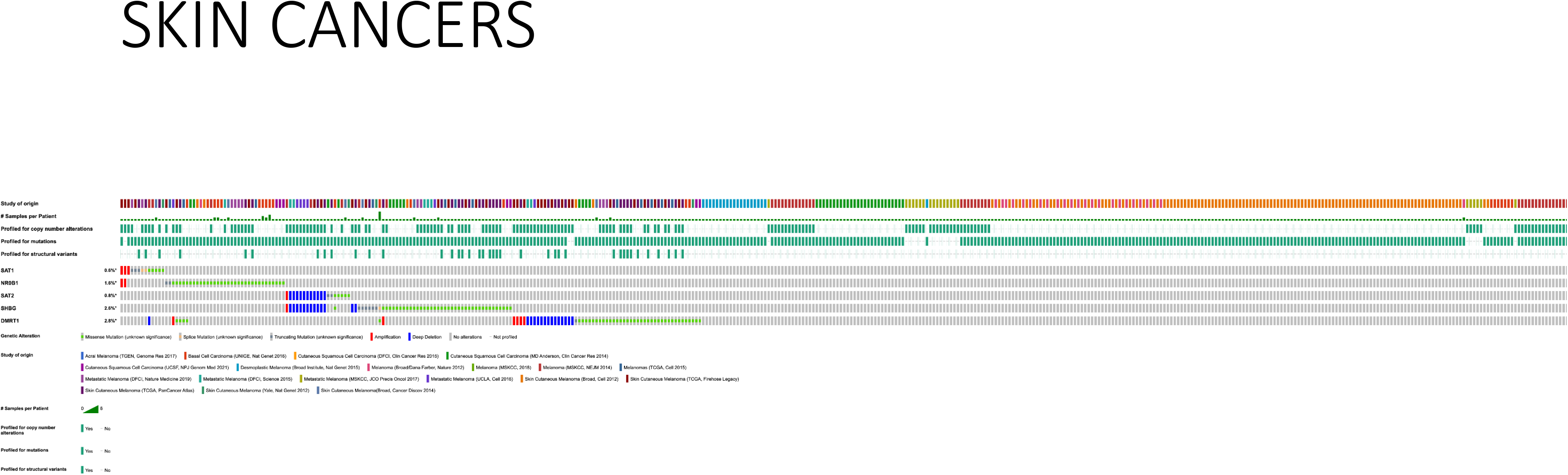
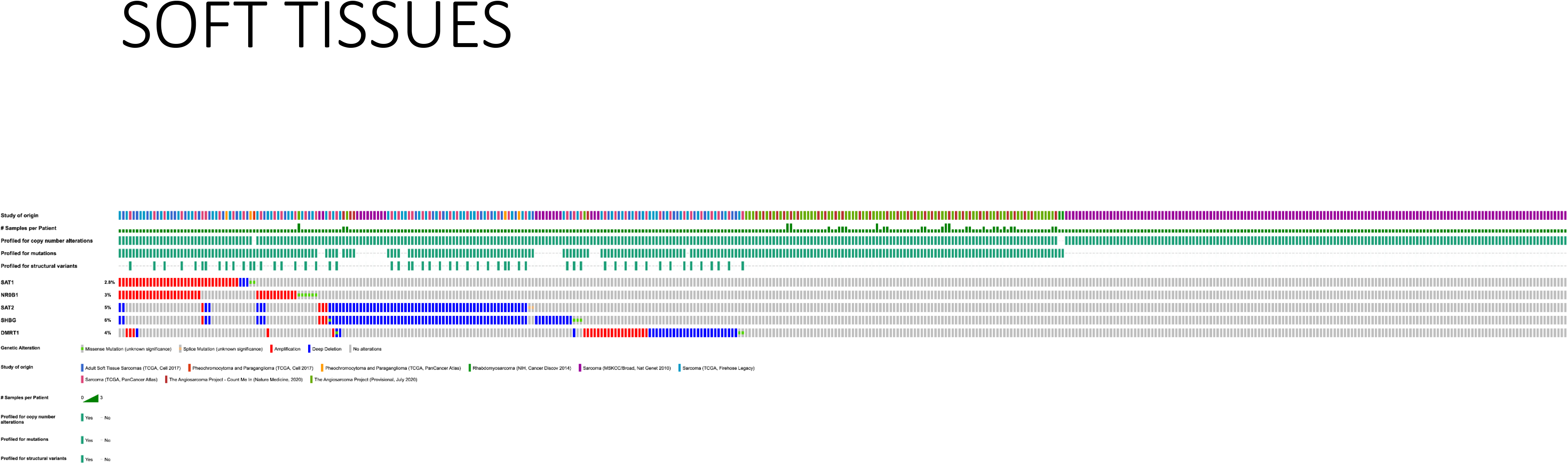
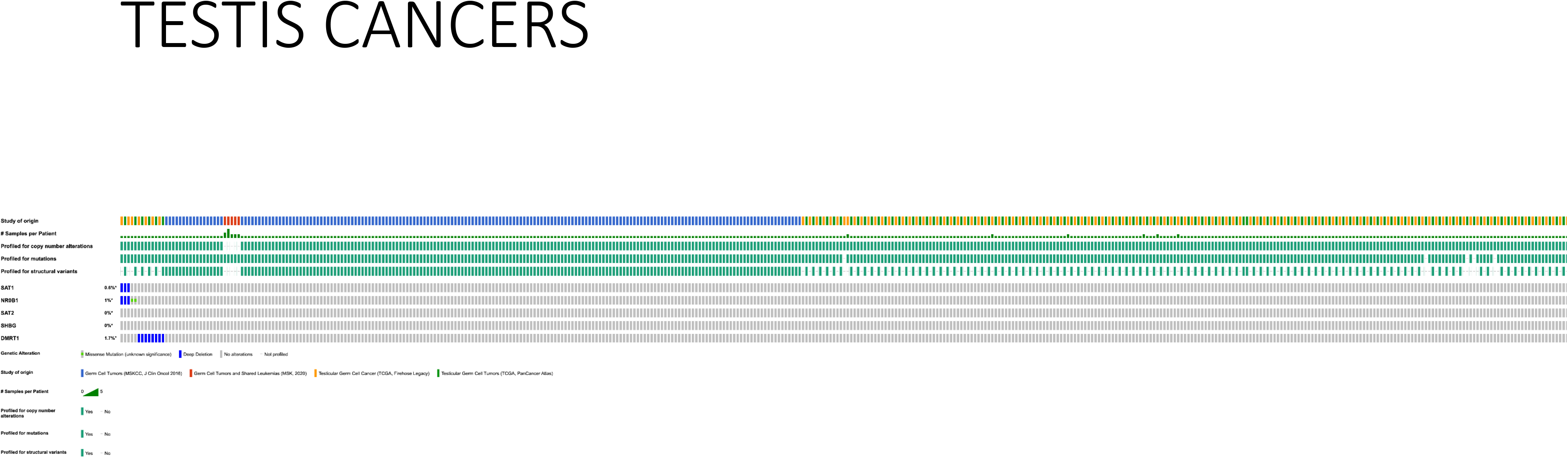
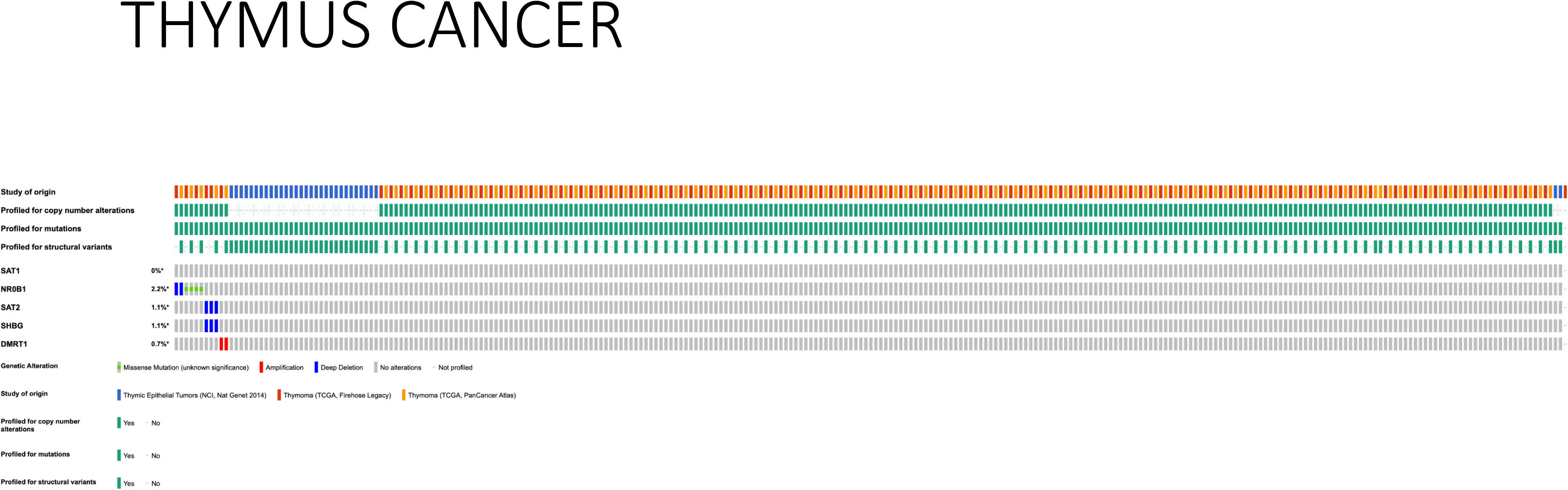
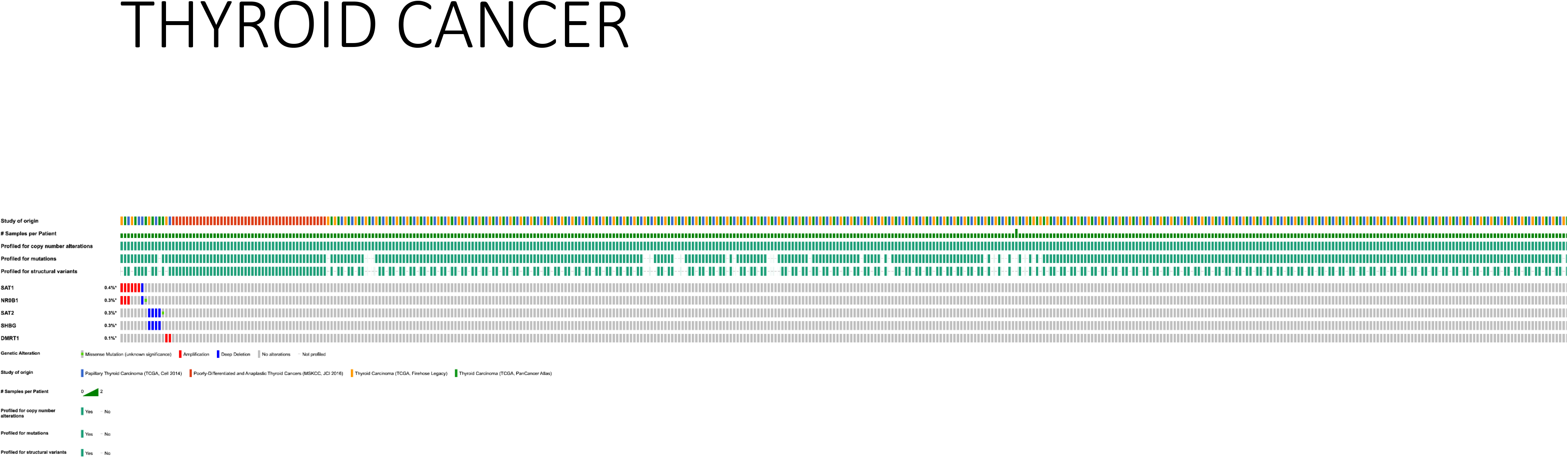
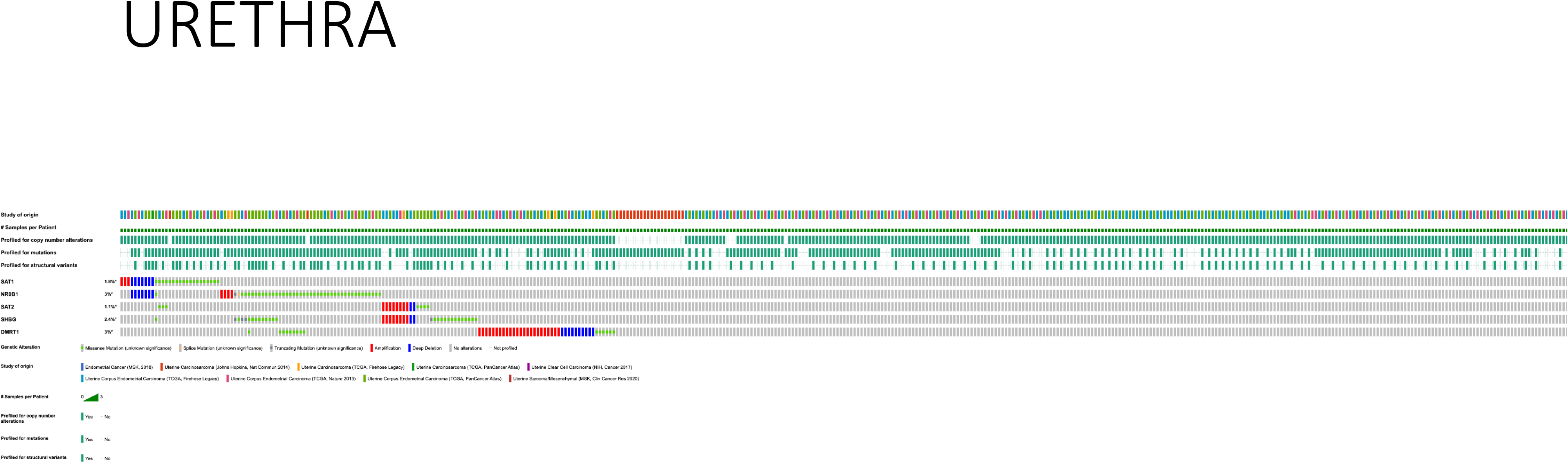

